# Determinants of Heart Rate Recovery and Heart Rate Variability in Lung Cancer Survivors Eligible for Long-Term Cure

**DOI:** 10.1101/508911

**Authors:** Duc Ha, Atul Malhotra, Andrew L. Ries, Wesley T. O’Neal, Mark M. Fuster

## Abstract

**Background:** Lung cancer survivors are at increased risk for autonomic dysfunction. We aimed to identify determinants of parasympathetic nervous system (PNS) function as reflected by heart rate recovery (HRR) and heart rate variability (HRV) in lung cancer survivors eligible for long-term cure.

**Methods:** We performed a cross-sectional study of consecutive lung cancer survivors who completed curative-intent therapy for stage I-IIIA ≥1 month previously. We tested a comprehensive list of variables related to baseline demographics, comorbidities, lung cancer characteristics, and physiological/functional measures using univariable and multivariable (MVA) linear regression analyses. We defined HRR as the difference in heart rate (HR) at 1-minute following and the end of the six-minute walk test (6MWT), and HRV the standard deviation of normal-to-normal R-R intervals (SDNN) and root-mean-square-of-successive-differences (rMSSD) from routine single 10-s electrocardiographs (ECGs).

**Results:** In 69 participants, the mean (standard deviation, SD) HRR was -10.6 (6.7) *beats.* In MVAs, significant independent determinants of HRR [β (95% confidence interval)] were: age [0.17 (0.04, 0,30) for each *year*] and HR change associated with the 6MWT [0.01 (0.007, 0.02) for each *beats/min.* In 41 participants who had ECGs available for HRV measurements, the mean (SD) SDNN and rMSSD were 19.1 (15.6) and rMSSD 18.2 (14.6) *ms,* respectively. In MVAs, significant determinants of HRV were: total lung capacity [0.01 (0.00, 0.02), p=0.047 for each *% predicted*] and HRR [-0.04 (-0.07, -0.003) for each *beat*] for natural logarithm (Ln-)SDNN; and [0.01 (0.00, 0.02)] and [-0.04 (-0.07, -0.01)] for Ln-rMSSD, respectively.

**Conclusions:** We measured determinants of HRR and HRV in lung cancer survivors eligible for long-term cure. HRR and/or HRV may be useful as indicators to stratify patients in interventional studies aimed at improving PNS function in lung cancer survivors, including through exercise training.

## Introduction

Lung cancer survivors are at risk for health impairments associated with the effects of aging, health behaviors, comorbidities, and/or lung cancer and its treatment [1]. Physiological evaluation in lung cancer is most commonly performed to measure or estimate peak oxygen consumption, an independent predictor of perioperative morbidity and mortality in patients being considered for major lung resection [2]. More recently, the utility of physiological evaluation outside of the preoperative context has been described, including in post-treatment lung cancer survivors to identify health impairments [3].

The autonomic nervous system (ANS) plays important physiological roles in the homeostasis of important organs, including heart and lungs [4]. Autonomic dysfunction (AD) as reflected by decreased parasympathetic tone and increased sympathetic tone has been reported in hypertension, hyperlipidemia, diabetes, heart failure, chronic obstructive pulmonary disease (COPD), and depression/anxiety [5]. Tobacco exposure [6] and antineoplastic therapy [7] can also lead to AD. Moreover, in a landmark study of patients undergoing exercise testing, AD was found to be associated with poor survival independent of standard cardiac risk factors [8]. The ANS therefore can be another domain of physiological evaluation that has diagnostic, predictive, and prognostic utility in lung cancer patients.

Heart rate recovery (HRR) following exercise testing is a marker of parasympathetic nervous system (PNS) function which is associated with clinical outcomes in various patient populations [8, 9]. Heart rate variability (HRV) is also a marker of PNS function [10] that can be more readily obtained from routine resting electrocardiographs (ECGs) and has reference values available for interpretation [11]. In recent years, HRV obtained from short-term single and repeated ECGs has been validated [12] and shown to be prognostic independent of cardiac risk factors [13] and in individuals without cardiovascular disease [11]. The identification of determinants of HRR and HRV may provide important insights into factors that could be modified to improve PNS function. In this study, we aimed to identify determinants of HRR and HRV in lung cancer survivors eligible for long-term cure. We hypothesized that HRR and HRV are inter-dependently associated, potentially through vagal activity, and can be used to assess PNS function in these patients.

## Methods

### Study Overview

We previously reported a cross-sectional study to characterize functional capacity and cancer-specific quality of life (QoL) in lung cancer survivor following curative-intent treatment [14]. In the present study, we enrolled additional participants to identify determinants of HRR and HRV and explore their inter-dependence. In brief, we identified eligible patients from a tumor board list of consecutive lung cancer cases managed at the VA San Diego Healthcare System (VASDHS). Between July 2016 and March 2018, we mailed information letters to eligible patients diagnosed with/managed for lung cancer at the VASDHS from October 2010 to February 2018 and followed up with a telephone call approximately one week later to gauge their interest. All enrolled participants provided written informed consent. This protocol was approved by the VASDHS Institutional Review Board (no. H150158).

### Participants

We enrolled lung cancer survivors who completed curative-intent lung cancer treatment, defined as lung cancer resection surgery, definitive radiation, or concurrent chemoradiation for stage I-IIIA disease ≥1 month previously (Figure 1). We excluded patients who were unable to perform exercise testing/HRR evaluation, due to severe dementia (n=2), bilateral below-knee amputation (n=2), or quadriplegia (n=1). For HRV analyses, we used outpatient ECGs obtained clinically within six months of enrollment, and excluded patients with atrial arrhythmias, atrial or ventricular pacing, or frequent premature atrial or ventricular contractions.

**Figure 1:**
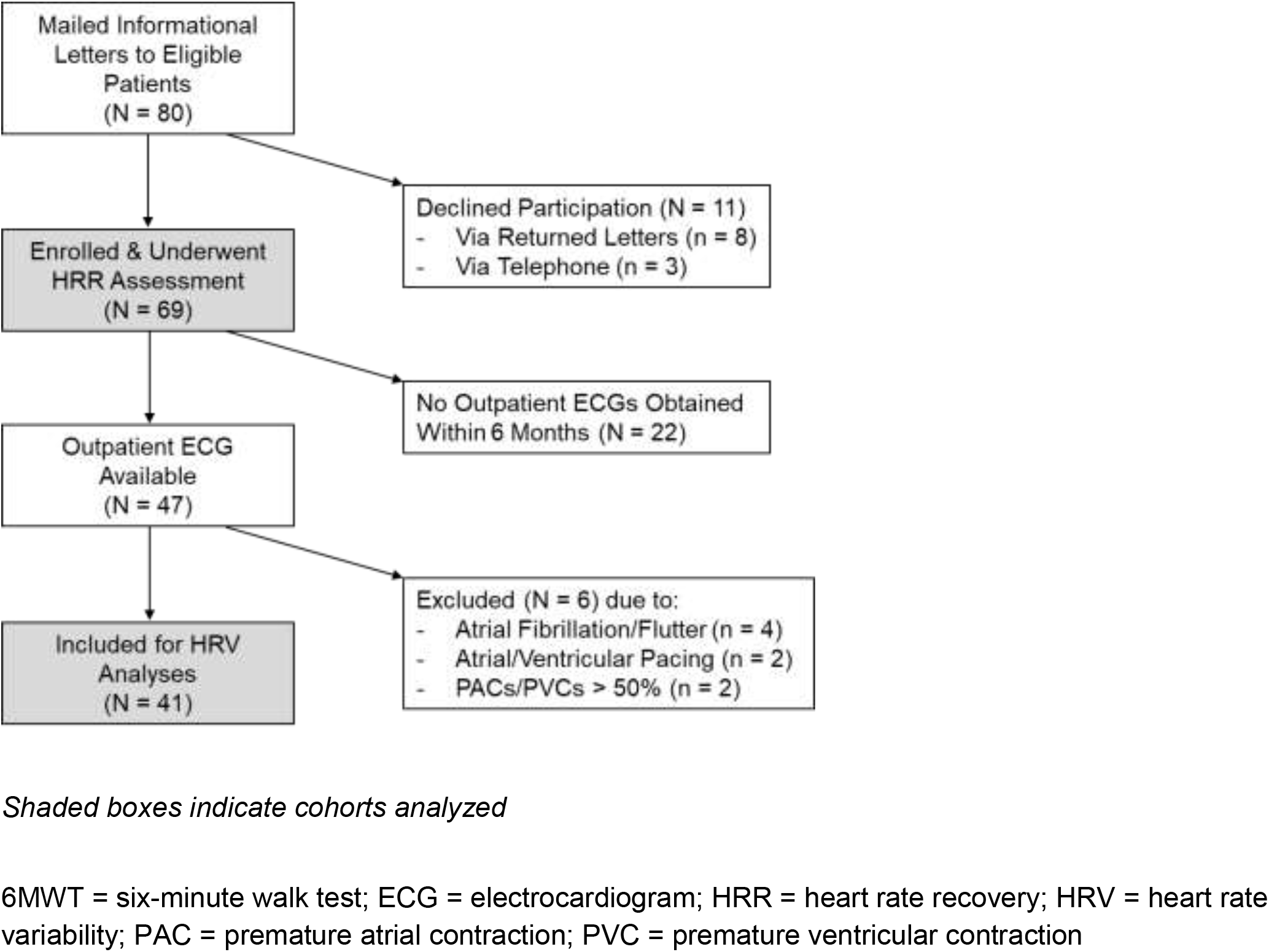
Flow Diagram of Included Participants

### Variables

We collected clinical characteristics which may be related to autonomic function including age, sex, ethnicity, smoking history, comorbidities (e.g. chronic cardiovascular and pulmonary diseases), medications (e.g. beta-blockers, inhibitors of the renin-angiotensin system), lung function [forced expiratory volume in 1 second (FEV_1_), total lung capacity (TLC), diffusion capacity of the lung for carbon monoxide (DL_CO_)], and echocardiographic findings where available. Lung cancer characteristics included histologic subtype, stage, and treatment.

### Heart Rate Recovery

We obtained HRR using the six-minute walk test (6MWT) as supported by existing literature [9, 15]. We performed the 6MWT according to the standard protocol at the VASDHS which follows the American Thoracic Society Pulmonary Function Standards Committee recommendations [16], modified to include HRR evaluation. In brief, we obtained pre-6MWT vital signs with the patient in the seated position. We then instructed participants to walk as far as possible for six minutes in a 130-ft (40-m) hallway. At the end of six minutes, we instructed participants to sit down for post-6MWT measures. We used a finger-probe pulse oximeter to obtain heart rate (HR) and oxygen (O_2_) saturation levels, and defined HRR as the difference, in *beats,* in HR at 1-minute following completion of the 6MWT and at the end of the 6MWT. Participants who had supplemental O_2_ prescribed used their own equipment at the same flow rate as their regular prescription. Practice tests and ECG monitoring were not performed, as per ATS recommendations [16].

### Heart Rate Variability

We assessed HRV using the standard deviation of normal-to-normal R-R intervals (SDNN) and root-mean-square of successive differences (rMSSD) from routine 12-lead 10-s ECG tracings, previously shown to correlate and agree well (Pearson’s *r* 0.85-0.86, Bland-Altman limits of agreement 0.08-0.10, and Cohen’s d 0.15-0.17 for rMSSD) with the gold-standard 240-300-s tracings [12]. We visually inspected all ECGs and excluded those with atrial arrhythmias or atrial or ventricular pacing. We additionally excluded premature atrial/ventricular contractions (PAC/PVCs) and the beats before and after them [11]; ECGs with >50% beats associated with PAC/PVCs were excluded as suggested by previous literature [11]. We measured normal-to-normal R-R intervals manually using electronic calipers in the GE^®^ MUSE editor software. All 6MWT, HRR, and HRV assessments were conducted by one observer blinded to all baseline characteristics and HRR or HRV measurements (DH).

### Statistical Analyses

We summarized descriptive statistics as means and standard deviations (SDs) for all continuous variables and as counts and percentages for categorical variables. Both HRR and HRV were recorded and analyzed as continuous variables. We interpreted the six-minute walk distance (6MWD) using reference equations in healthy adults [17], HRR using a cutoff of ≤12-*beat* decrease to indicate impairment [8, 9], and SDNN and rMSSD using reference values for stage I-II non-small cell lung cancer (NSCLC) patients [18]. We also used reference values provided in the literature to compare HRV measures between our cohort and historical controls.

We transformed SDNN and rMSSD into normal distribution using natural logarithms as supported by previous literature [18]. We used correlation coefficients, and univariable (UVA) and multivariable (MVA) linear regressions to assess and analyze the relationship between baseline characteristics and HRR and HRV. We performed MVAs using stepwise, backward selection modeling starting with all baseline characteristics with p <0.20 in UVAs, and used regression coefficients (β), 95% confidence intervals (CIs), and coefficients of determination (R^2^ and partial R^2^) for interpretation. Statistical significance was defined as p <0.05 in two-tailed tests. All data were entered and managed using REDCap electronic data capture tools hosted at the University of California San Diego Clinical and Translational Research Institute [19]. IBM^®^ SPSS^®^ Statistics software version 23.0 was used for all analyses.

## Results

### Participants

We enrolled 69 participants who completed 6MWT and HRR evaluation; their baseline clinical characteristics are described in Table 1. Most participants were white males who were current or former smokers and underwent either lung cancer resection surgery or definitive radiation for treatment.

**Table 1:** Participant Characteristics

| Participant Characteristic<br>(N=69, VASDHS, 2016-2018) | Value |
| --- | --- |
| Age, mean (SD) | 71.0 (8.4) |
| White race, n (%) | 63 (91) |
| Male sex, n (%) | 66 (96) |
| BMI, kg/m<sup>2</sup>, mean (SD) | 26.5 (4.8) |
| Smoking history, n (%) |  |
| Current | 22 (32) |
| Former | 41 (59) |
| Never | 6 (9) |
| Pack years, mean (SD) | 52.8 (32.2) |
| Comorbidities, n (%) |  |
| Hypertension | 57 (83) |
| Hyperlipidemia | 57 (83) |
| Diabetes | 18 (26) |
| Atrial arrhythmia* | 17 (25) |
| CAD | 25 (36) |
| HF<sup>†</sup> | 17 (25) |
| COPD/asthma | 51 (74) |
| OSA | 16 (23) |
| Anxiety/Depression/PTSD | 21 (30) |
| Other cancers | 29 (42) |
| Medications, n (%) |  |
| Beta-blockers | 31 (45) |
| ACE-I/ARB | 27 (39) |
| Pulmonary function, mean (SD) |  |
| FEV<sub>1</sub>/FVC, % | 59.9 (14.9) |
| FEV<sub>1</sub>, % *predicted* | 71.1 (25.2) |
| TLC, % *predicted* ** | 110.9 (22.3) |
| DL<sub>CO</sub>, % *predicted* | 78.6 (25.6) |
| Ventilatory defects<sup>‡</sup>, n (%) |  |
| Obstructive | 51 (74) |
| Restrictive | 3 (4) |
| DL<sub>CO</sub> limitation | 38 (55) |
| Lung cancer characteristics |  |
| Clinical stage, n (%) |  |
| I | 55 (80) |
| II | 4 (6) |
| IIIA | 10 (15) |
| Histologic subtype, n (%) |  |
| Adenocarcinoma | 33 (48) |
| Squamous cell carcinoma | 17 (25) |
| Presumed | 14 (20) |
| Primary treatment, n (%) |  |
| Surgical resection | 34 (49) |
| Definitive radiation | 25 (36) |
| Chemoradiation | 10 (15) |

\*Any history of, including paroxysmal atrial fibrillation or flutter, treated with medical therapy or ablation.
\*\*Data available in 58 participants.
<sup>†</sup>Defined as ejection fraction <50% or clinical documentation of systolic heart failure.
<sup>‡</sup>Defined as $FEV_1/FVC < 70\%$ for obstructive and $TLC \% \text{ predicted} < 80$ for restrictive defects, and $DL_{CO} \% \text{ predicted} < 80$ for $DL_{CO}$ limitation.
ACE-I = angiotensin converting enzyme inhibitor; ARB = angiotensin II receptor blocker; BMI = body-mass index; CAD = coronary artery disease; COPD = chronic obstructive pulmonary disease; $DL_{CO}$ = diffusion capacity of the lung for carbon monoxide; $FEV_1$ = forced expiratory volume in 1 second; FVC = forced vital capacity; HF = heart failure; OSA = obstructive sleep apnea; PTSD = posttraumatic stress disorder; SD = standard deviation; TLC = total lung capacity

### Determinants of HRR

The mean (SD) 6MWD for all participants was 342 (123) *m*, 66 (24) % predicted, with 40 participants (58%) having impaired functional capacity as defined by standard equations for normal healthy adults [17]. Following the 6MWT, the mean (SD) HRR was -10.6 (6.7) *beats*; 45 participants (65%) had impaired HRR as defined by a cutoff of ≤12-*beat* decrease (Table 2) [8, 9].

**Table 2:** Physiological Measures

| 6MWT-Associated Measures (N=69) | Values |
| --- | --- |
| Pre-6MWT Vital Signs |  |
| HR, *beats/min* | 74.0 (13.7) |
| SBP, *mmHg* | 130.2 (18.4) |
| DBP, *mmHg* | 78.3 (11.9) |
| O<sub>2</sub> saturation, % | 96.6 (2.0) |
| Physiological change, mean (SD) |  |
| HR, *beats/min* | +20.4 (11.3) |
| SBP, *mmHg* | +13.5 (17.2) |
| DBP, *mmHg* | +1.9 (6.9) |
| O<sub>2</sub> saturation, % | -3.1 (4.5) |
| Borg dyspnea score, mean (SD) |  |
| Pre-6MWT | 0.84 (1.4) |
| Post-6MWT | 3.8 (2.6) |
| Change | +2.9 (2.4) |
| Functional EC |  |
| 6MWD, *m*, mean (SD) | 341.5 (123.4) |
| 6MWD, % *predicted*, mean (SD) | 65.6 (24.3) |
| Impaired EC (6MWD < LLN), n (%) | 40 (58) |
| Heart rate recovery* |  |
| HRR, *beats*, mean (SD) | -10.6 (6.7) |
| Impaired HRR, n (%) | 45 (65) |
| Heart rate variability** (N=41) |  |
| SDNN, *ms*, mean (SD) | 19.1 (15.6) |
| Impaired SDNN, n (%) | 24 (59) |
| rMSSD, *ms*, mean (SD) | 18.2 (14.6) |
| Impaired rMSSD, n (%) | 24 (59) |

\*Data obtained post-treatment in all (n=69) participants.
\*\*Data were obtained from pre-treatment (n=14) and post-treatment (n=27) electrocardiographs.
6MWD = six-minute walk distance; 6MWT = six-minute walk test; DBP = diastolic blood pressure; EC = exercise capacity; HR = heart rate; HRR = heart rate recovery; LLN = lower-limit of normal; O<sub>2</sub> = oxygen; SBP = systolic blood pressure; SD = standard deviation

In UVA, hyperlipidemia, atrial arrhythmia, pre-6MWT HR, HR change associated with the 6MWT, and 6MWD were associated with HRR (Table 3); lung cancer histologic subtype, stage, and primary treatment modality (surgical resection, definitive radiation, or chemoradiation) were not associated with HRR (E-Table 2). In MVAs starting with all baseline characteristics with p <0.20 in UVAs, in a final model (Table 4Ai) that also included hyperlipidemia and pre-6MWT, age and HR change associated with the 6MWT were significant independent determinants of HRR (Figure 2Ai-ii & E-Figure 1A-B). When patients with a history of paroxysmal, persistent, or permanent atrial arrhythmia managed with medical and/or ablation therapy were excluded (UVA results are shown in E-Table 1), similar results were obtained; heart failure with reduced ejection fraction was additionally found to be a significant determinant of HRR (Table 4Aii).

**Table 3:**
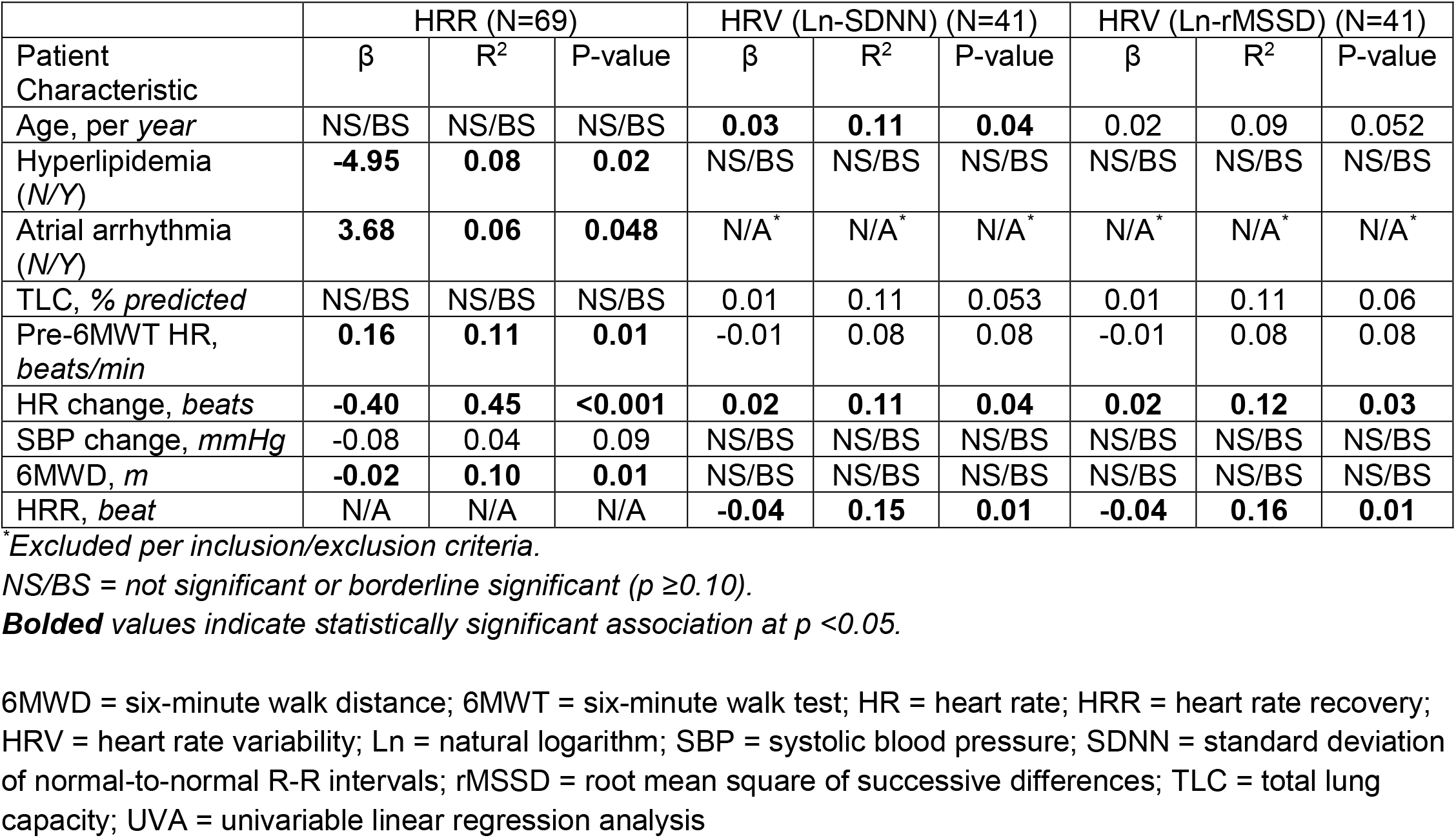
UVA – Significant/Borderline Determinants of HRR and HRV

**Table 4Ai:**
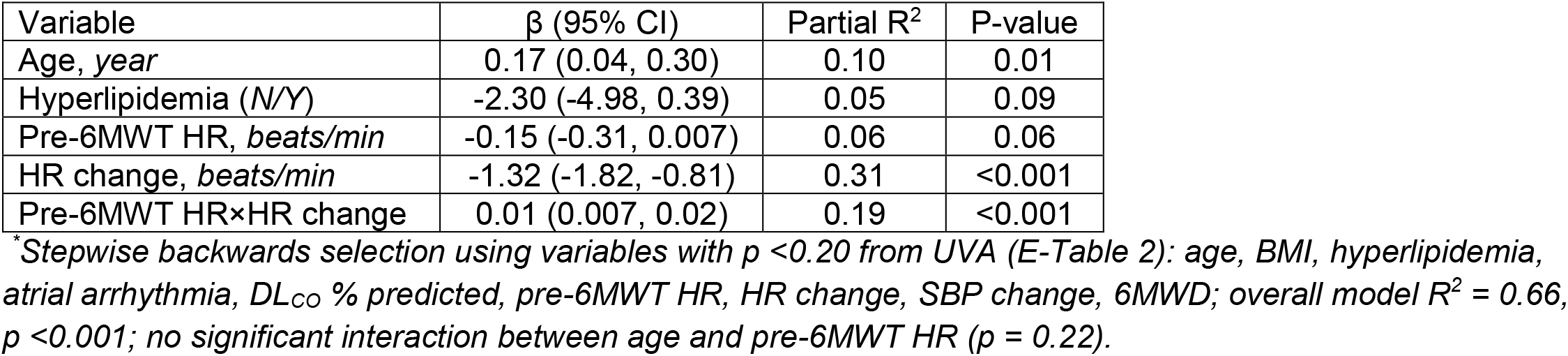
MVA* – Independent Determinants of HRR (N=69)

| Variable | $\beta$ (95% CI) | Partial R<sup>2</sup> | P-value |
| --- | --- | --- | --- |
| Age, year | 0.17 (0.04, 0.30) | 0.10 | 0.01 |
| Hyperlipidemia (N/Y) | -2.30 (-4.98, 0.39) | 0.05 | 0.09 |
| Pre-6MWT HR, beats/min | -0.15 (-0.31, 0.007) | 0.06 | 0.06 |
| HR change, beats/min | -1.32 (-1.82, -0.81) | 0.31 | <0.001 |
| Pre-6MWT HR×HR change | 0.01 (0.007, 0.02) | 0.19 | <0.001 |

\*Stepwise backwards selection using variables with $p < 0.20$ from UVA (E-Table 2): age, BMI, hyperlipidemia, atrial arrhythmia, DL<sub>CO</sub> % predicted, pre-6MWT HR, HR change, SBP change, 6MWD; overall model $R^2 = 0.66$ , $p < 0.001$ ; no significant interaction between age and pre-6MWT HR ( $p = 0.22$ ).

**Figure 2A:**
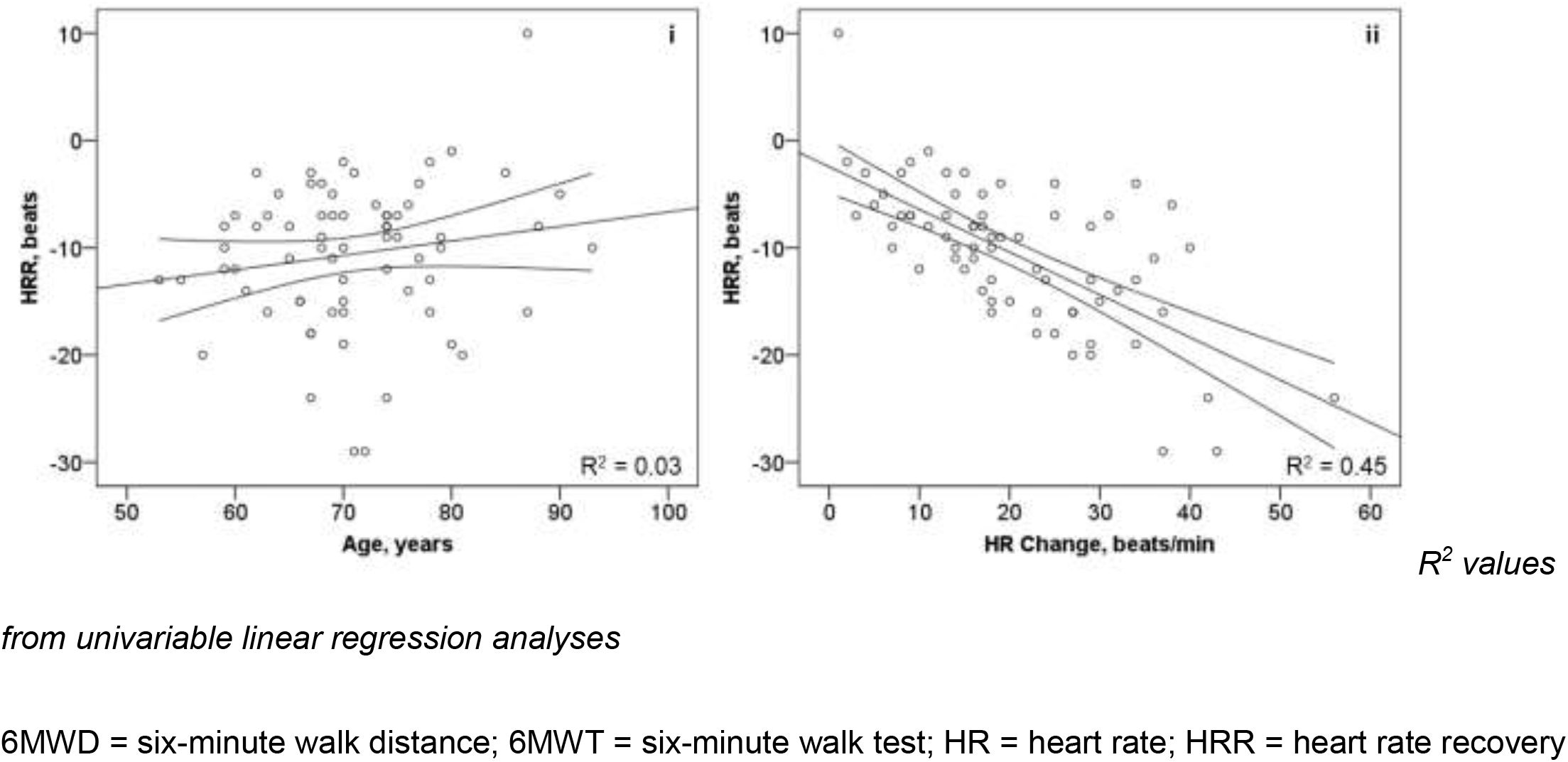
Significant Independent Determinants of HRR

**Table 4Aii:**
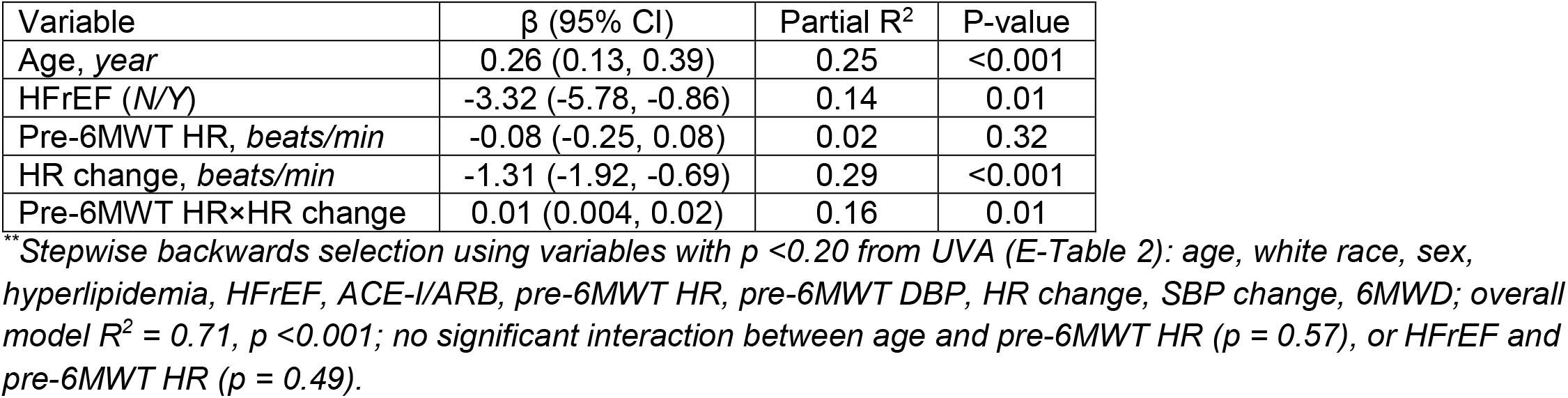
MVA** – Independent Determinants of HRR (excluding atrial arrhythmia) (N=52)

### Determinants of HRV

Forty-one participants had ECGs obtained clinically within 6 months of enrollment and met criteria for HRV evaluation (Figure 1). There were no significant differences in clinical characteristics between those with and without ECGs available except for the change in diastolic blood pressure associated with the 6MWT (E-Table 2). The mean (SD) SDNN and rMSSD were 19.1 (15.6) and 18.2 (14.6) *ms,* respectively, and their natural logarithm transformed values 2.72 (0.64) and 2.69 (0.63).

Twenty-four participants (59%) had impaired SDNN and rMSSD, defined as <the mean reference values derived from single 10-s ECGs for stage I-II NSCLC patients [18]. There was no significant difference in SDNN (p = 0.33) or rMSSD (p = 0.27) between our cohort and historical stage I-II NSCLC patients [18]. Compared to individuals without cardiovascular disease included in the Multi-Ethnic Study of Atherosclerosis [11], lung cancer survivors included in our cohort had a mean difference of -5.04 *ms* (95% CI -9.96, -0.12, p = 0.045) in SDNN and -9.08 *ms* (95% CI -13.7, -4.47, p <0.001) in rMSSD.

In UVA (E-Table 3), clinical characteristics significantly/borderline associated with HRV were age, TLC *% predicted,* pre-6MWT HR, HR change associated with the 6MWT, and HRR following the 6MWT (Table 3). Exploratory UVA in 27 participants with ECGs obtained post-lung cancer treatment showed no significant association between primary treatment modality and HRV (p = 0.36 for Ln-SDNN, and p = 0.37 for Ln-rMSSD). In MVA (Table 4B) in a model that also contained age, TLC *% predicted* and HRR (Figure 2Bi-ii, E-Figure 1A-B) were significant independent determinants of HRV.

**Figure 2B:**
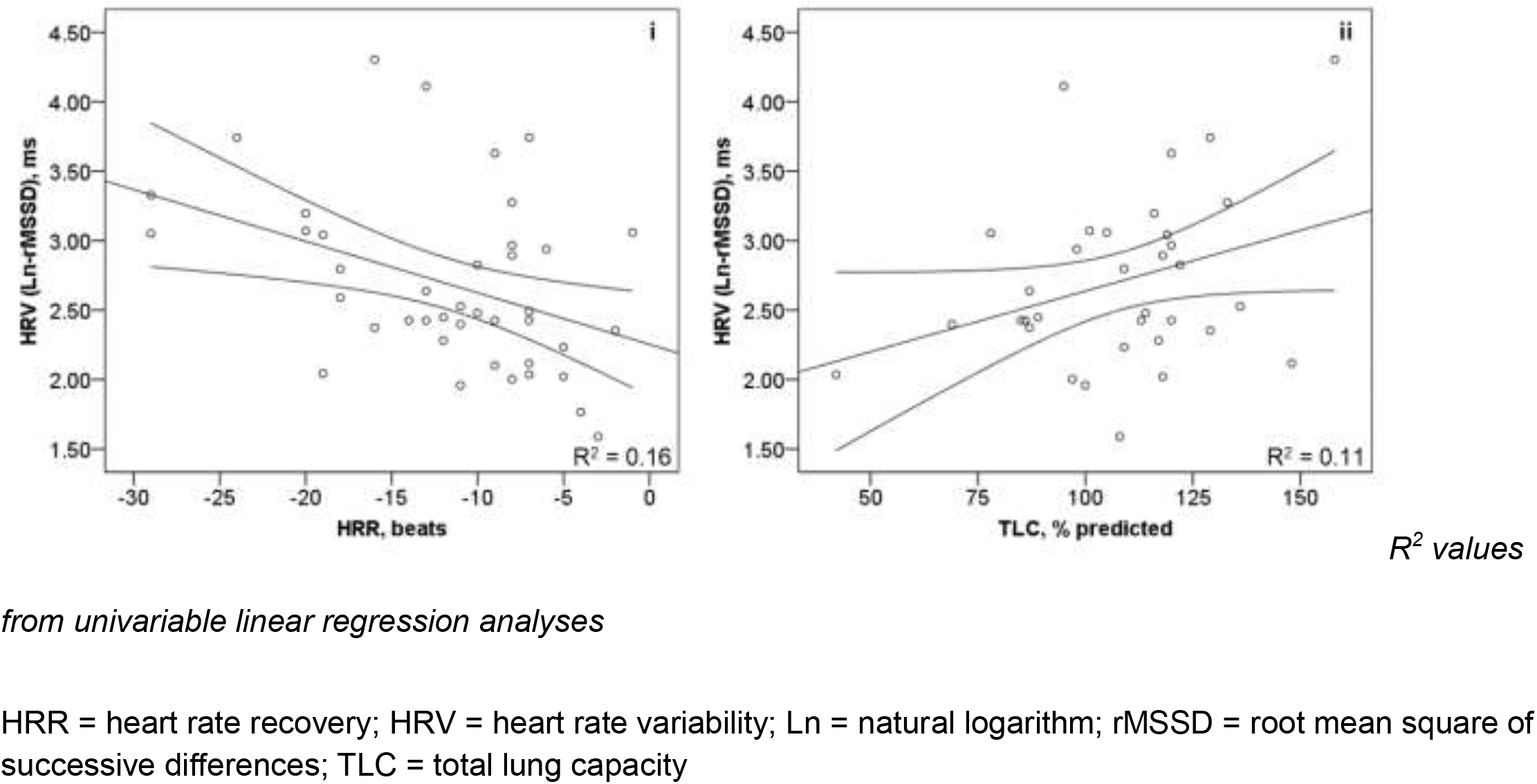
Significant Independent Determinants of HRV (Ln-rMSSD)

**Table 4B:** MVA – Independent Determinants of HRV (N=41)

| Variable | Ln-SDNN<sup>a</sup> |  |  | Ln-rMSSD<sup>b</sup> |  |  |
| --- | --- | --- | --- | --- | --- | --- |
| | $\beta$ (95% CI) | Partial R<sup>2</sup> | P-value | $\beta$ (95% CI) | Partial R<sup>2</sup> | P-value |
| Age, year | 0.02 (-0.01, 0.04) | 0.07 | 0.14 | 0.01 (-0.01, 0.04) | 0.05 | 0.199 |
| TLC, % predicted | 0.01 (0.00, 0.02) | 0.13 | 0.047 | 0.01 (0.00, 0.02) | 0.13 | 0.047 |
| HRR, beat | -0.04 (-0.07, -0.003) | 0.14 | 0.03 | -0.04 (-0.07, -0.01) | 0.16 | 0.02 |

<sup>a</sup>Stepwise backwards selection using variables with $p < 0.20$ from UVA (E-Table 3): age, TLC % predicted, Pre-6MWT HR, HR change, O<sub>2</sub> sat change, HRR; overall model $R^2 = 0.31$ , $P = 0.01$ ; no significant interaction between TLC % predicted and HRR ( $p = 0.46$ ).
<sup>b</sup>Stepwise backwards selection using variables with $p < 0.20$ from UVA (E-Table 3): age, TLC % predicted, Pre-6MWT HR, HR change, O<sub>2</sub> sat change, HRR; overall model $R^2 = 0.30$ , $P = 0.01$ ; no significant interaction between TLC % predicted and HRR ( $p = 0.41$ ).
6MWD = six-minute walk distance; 6MWT = six-minute walk test; ACE-I = angiotensin converting enzyme inhibitor; ARB = angiotensin II receptor blocker; BMI = body-mass index; CI = confidence interval; DBP = diastolic blood pressure; DL<sub>CO</sub> = diffusion capacity of the lung for carbon monoxide; HFrEF = heart failure with reduced ejection fraction; HR = heart rate; HRR = heart rate recovery; HRV = heart rate variability; Ln = natural logarithm; MVA = multivariable linear regression analysis; SBP = systolic blood pressure; SDNN = standard deviation of normal-to-normal R-R intervals; rMSSD = root mean square of successive differences; TLC = total lung capacity; UVA = univariable linear regression analysis

### Characterization of HRR-HRV Inter-Dependence

Heart rate recovery following the 6MWT correlated moderately-well with HRV (Spearman’s ρ -0.38, p = 0.01 for SDNN and -0.41, p = 0.008 for rMSSD, respectively). Impaired SDNN/rMSSD was concordant with HRR in 69% of cases. Overall, the mean HRR for participants with normal compared to impaired SDNN and rMSSD, respectively were -14.4 vs. -9.7 *beats* (p = 0.04) and -15.6 vs. -9.8 *beats* (p = 0.04) (E-Figure 2A-B).

## Discussion

In lung cancer survivors eligible for long-term cure, we measured determinants of HRR and HRV and found impairments in 65% in 59% of patients, respectively. In addition, HRV and HRR were inter-dependently associated, supporting their utility to assess PNS function in the lung cancer population. To the best of our knowledge, our study is the first to measure determinants of HRR and HRV in lung cancer survivors eligible for long-term cure and characterize their inter-relationship. Similar to a previous study [20], we report a moderate correlation (correlation coefficient 0.3-0.7) between SDNN/rMSSD and HRR.

Lung cancer survivors have various etiologies implicated with AD, including due to aging [21], tobacco exposure [22], prevalence of comorbidities including COPD [23], heart failure [24], and diabetes [25], and chemotherapy treatment [26]. In our multivariable models, HRR explains approximately 15% of the variances in HRV, suggesting caution against exchanging HRR for HRV to assess PNS function. Notably, HRR can vary depending on cardiopulmonary fitness, exertional levels achieved, and changes in HR associated with exercise testing [27], including in cancer survivors [28]. Variations in HRR may also be related to chronotropic incompetence [29] which may be present in some participants included. In addition, HRV from 10-s ECGs have increased agreement with the gold-standard 240-300-s tracings when repeated measurements are obtained (up to 3 times) [12]; we obtained HRV from single 10-s ECGs which may additionally explain some of the variances between HRR and HRV.

Similar to previous studies, we found HRR to be associated with age, resting and peak HR, exercise capacity, but not with beta-blocker or renin-angiotensin system inhibitor use [8, 25]. Like a previously study involving 154 COPD patients [30], we found no association between HRV and functional capacity; this is in contrast to another study [25] which reported a significant independent association between HRV as reflected by SDNN and rMSSD and exercise capacity in 1,060 patients with coronary artery disease (CAD) with and without type 2 diabetes (DM). Differences in comorbidities (all/predominantly COPD vs. CAD with/without DM) and sample size may explain these contrasting findings. We found a positive association in TLC and HRV, similar to a previous report involving 30 patients with stable COPD [31]. This finding may have important implications in future studies aimed at assessing HRV in the lung cancer population in which COPD is highly prevalent [32] and could reflect alterations in vagal and pulmonary stretch receptor activities associated with chronic hyperinflation [33]. Unlike existing literature [34, 35], age was not associated with HRV in multivariable analyses, possibly due to the inclusion of age in TLC *% predicted* which adjusts for age in our models or a small sample size.

Our study also suggests that on average, those with normal HRV have an average HRR of 14-to 16-beat decreases which are significantly higher than those with impaired HRV. Previous studies in the cardiovascular and chronic lung disease populations have suggested cutoff ranges of 12- to 18-*beat* decreases in HRR following exercise testing to predict outcomes [9]. Based on our data, a HRR cutoff of -14 to -16 *beats* may be a useful indicator of AD in lung cancer survivors eligible for long-term cure, similar to previous studies involving HRR following the 6MWT to predict acute exacerbations in COPD [36] and clinical worsening in pulmonary arterial hypertension [15] patients.

Traditionally, evaluation of exercise performance focuses on muscle and cardiovascular function. The nervous system is often under-recognized despite having important physiological bases: somatic innervations facilitate voluntary motor control, the sympathetic nervous system activates a “fight-or-flight” response at the beginning of exercise, and parasympathetic system a “rest-and-digest” state in recovery. While HRR is associated with exercise capacity partly due to the nature of the test (exercise is needed to assess recovery), a blunted HRR may be more closely related to physical inactivity than comorbidities according to one study [25]. Therefore, the ANS is another domain of fitness that may be useful in the lung cancer population. We previously reported associations between impaired HRR and perioperative cardiopulmonary complications following lung cancer resection surgery [37], and survival in patients undergoing stereotactic body radiotherapy for early-stage lung cancer [38]. Others have also reported an association between HRV and survival in cancer patients in a systematic review and meta-analysis [39]. Together, these data provide supporting evidence on the importance of the PNS in the lung cancer population.

Exercise has been shown to be effective in improving QoL and function in cancer survivors [40]. However, the evidence of effectiveness is not as consistent in lung cancer survivors compared to other cancers [41], possibly due to unique characteristics in lung cancer as discussed above. Physiological evaluation in lung cancer survivors may help identify patients at risk for clinical worsening who may benefit from additional health services (e.g. rehabilitation and/or exercise programs) to improve health. Physiological measures that are readily available may help identify at-risk individuals, monitor their health changes, and evaluate the effectiveness of health interventions. HRR and HRV from 10-s ECGs are relatively easy to obtain, have been demonstrated to be responsive to exercise training [42, 43], and therefore may have such utility in lung cancer patients. Unlike HRR, HRV can be obtained at rest and therefore is not subjected to variations in patient effort associated with exercise testing.

Our study has limitations. First, our small sample size may not be adequately powered to detect significant associations between lung cancer-specific characteristics including stage and treatment modality and the ANS; however a previous analysis of 133 NSCLC patients reported that HRV on 10-s ECGs were lower in stage I-II compared to stage III-IV NSCLC [mean (SD) rMSSD = 15.6 (11.5) vs 20.3 (23.5) *ms,* respectively, p = 0.01] [18], suggesting lung cancer-specific effects on the ANS. Second, we did include other factors including ratings of perceived exertion (RPE) during the 6MWT which may limit our interpretation of the determinants of HRR; however the change in HR during exercise testing has been shown to correlate very well (correlation coefficient 0.74) with RPE [44] and therefore could be an indirect measure. Third, we did not measure HRV using the gold-standard 240-300-s tracings; however HRV measures from single 10-s ECGs have been shown to correlate and agree very well with the gold-standard as analyzed by correlation coefficients (*r* = 0.758 − 0.764 and 0.853 − 0.862 for SDNN and rMSSD, respectively), Bland-Altman 95% limits of agreements (bias = 0.398 − 0.416 and 0.079 − 0.096), and Cohen’s d statistics (d = 0.855 − 0.894 and 0.150 − 0.171) [12], and is prognostic in the elderly patient population [13]. Fourth, we did not assess other measures of ANS function including baroreflex sensitivity, muscle sympathetic nerve activity, or plasma catecholamines, and cannot give insights into the pathophysiological mechanisms relating AD in lung cancer survivors due to the descriptive nature of our study and the absence of related clinical outcomes, limiting definitive conclusions. Fifth, the single-institutional nature involving a predominantly white male veteran population and/or referral and survivor bias may limit the generalizability of our findings.

The strengths of our study include a thorough list of comorbidities and potential confounders relevant in the lung cancer population including tobacco exposure history and lung function. Also, all baseline characteristics were collected and verified by a board-certified physician to maximize accuracy, and all physiological assessments including the 6MWT, HRR, and HRV measurements were performed by one observer, limiting inter-operator variability. In addition, we performed thorough analyses to identify determinants of HRR and HRV, facilitating interpretation in the clinical setting. Last, we detected a moderate correlation/association between HRR and HRV, both of which are traditionally thought to reflect predominantly PNS function and therefore provided supporting evidence for their physiological basis in the lung cancer population.

We measured determinants of HRR and HRV in lung cancer survivors eligible for long-term cure. We conclude that HRR and HRV are inter-dependent measures of PNS function. These measures are relatively easy to obtain and may have diagnostic, predictive, and prognostic value. Studies aimed at improving health in lung cancer survivors, including through exercise training, may consider these measures for stratification and/or as physiological outcomes.

## Abbreviations

6MWD: six-minute walk distance
6MWT: six-minute walk test
AD: autonomic dysfunction
ANS: autonomic nervous system
ATS: American Thoracic Society
CAD: coronary artery disease
CI: confidence interval
COPD: chronic obstructive pulmonary disease
DL_CO_: diffusion capacity of the lung for carbon monoxide
DM: diabetes mellitus
ECG: electrocardiograph
FEV_1_: forced expiratory volume in 1 second
HR: heart rate
HRR: heart rate recovery
HRV: heart rate variability
MVA: multivariable linear regression analysis
NSCLC: non-small cell lung cancer
O_2_: oxygen
PAC: premature atrial contraction
PNS: parasympathetic nervous system
PVC: premature ventricular contraction
QoL: quality of life
rMSSD: root mean square of successive differences
RPE: ratings of perceived exertion
SDNN: standard deviation of normal-to-normal R-R intervals
TLC: total lung capacity
UVA: univariable linear regression analysis
VASDHS: VA San Diego Healthcare System

