## Supplemental Data for "Determinants of Heart Rate Recovery and Heart Rate Variability in Lung Cancer Survivors Eligible for Long-Term Cure"

**Supplementary Data**

E-Figure 1: Significant Independent Determinants of HRV (Ln-SDNN)

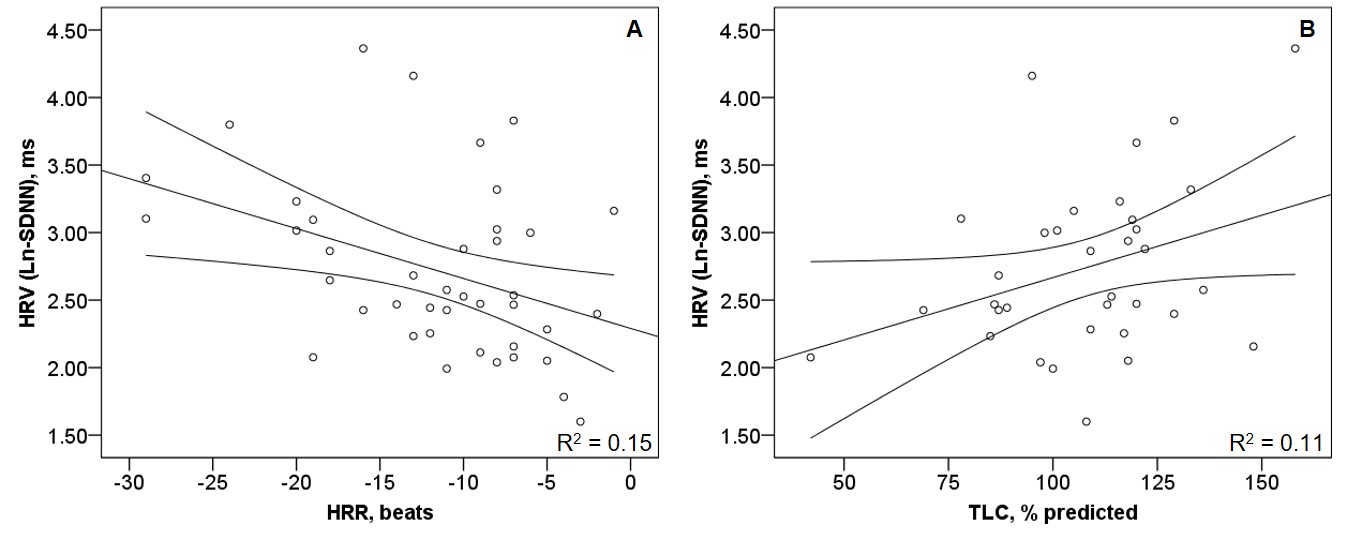

*R^2^ values from univariable linear regression analyses*

HRR = heart rate recovery; HRV = heart rate variability; Ln = natural logarithm; SDNN = standard deviation of normal-to-normal R-R intervals; TLC = total lung capacity

E-Figure 2: Differences in HRR between Impaired and Normal HRV

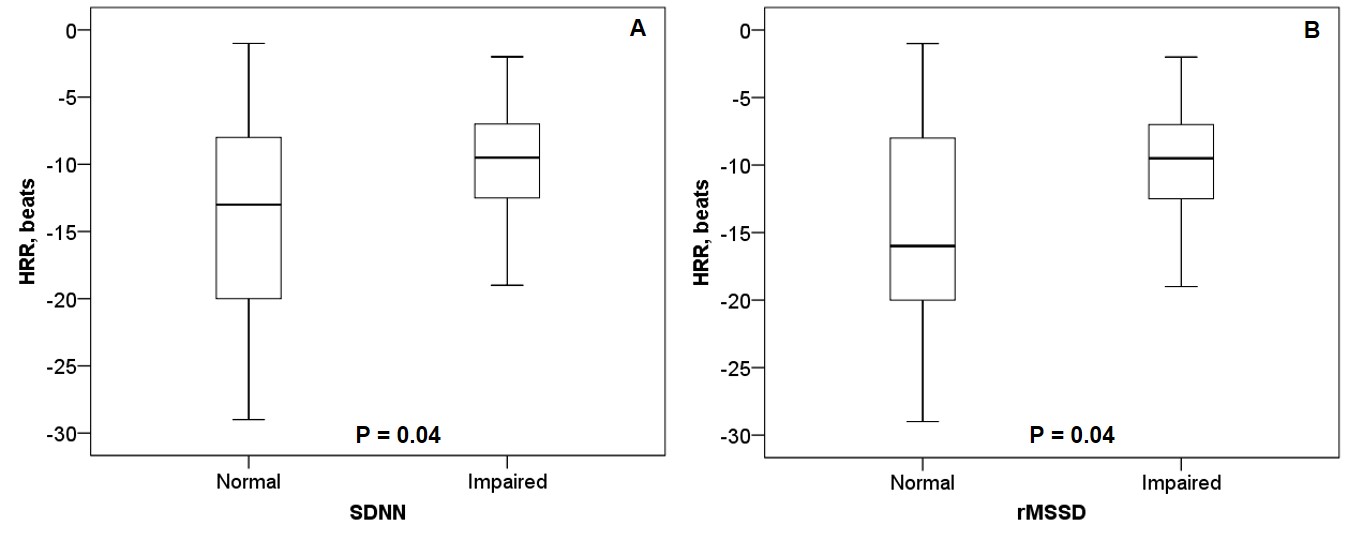

*P-values from independent sample t-tests (equal variances not assumed as determined by Levene’s test for equality of variances)*

HRR = heart rate recovery; HRV = heart rate variability; SDNN = standard deviation of normal-to-normal R-R intervals; rMSSD = root mean square of successive differences

E-Table 1: UVA – Determinants of HRR

|  | All Participants (N=69) | | | Excluding Atrial Arrhythmia (N=52) | | |
| --- | --- | --- | --- | --- | --- | --- |
| Patient Characteristic | β (95% CI) | R^2^ | P-value | β (95% CI) | R^2^ | P-value |
| Age, per year | 0.14 (-0.06, 0.33) | 0.03 | 0.17 | 0.16 (-0.04, 0.37 | 0.05 | 0.12 |
| White race (N/Y) | -2.47 (-8.18, 3.24) | 0.01 | 0.39 | -4.57 (-10.3, 1.18) | 0.05 | 0.12 |
| Sex (F/M) | -3.92 (-11.8, 3.95) | 0.02 | 0.32 | -4.95 (-12.3, 2.37) | 0.04 | 0.18 |
| BMI, per kg/m^2^ | -0.24 (-0.58, 0.09 | 0.03 | 0.15 | -0.16 (-0.52, 0.19) | 0.02 | 0.36 |
| Pack years, each | -0.001 (-0.05, -0.05) | 0.00 | 0.97 | 0.02 (-0.04, 0.07) | 0.01 | 0.48 |
| Hypertension (N/Y) | 0.70 (-3.56, 4.97) | 0.002 | 0.74 | -0.76 (-5.01, 3.49) | 0.003 | 0.72 |
| **Hyperlipidemia (N/Y)** | **-4.95 (-9.04, -0.85)** | **0.08** | **0.02** | **-6.30 (-10.2, -2.44)** | **0.18** | **<0.01** |
| DM (N/Y) | -2.36 (-6.00, 1.23) | 0.02 | 0.20 | -2.24 (-6.44, 1.97) | 0.02 | 0.29 |
| **Atrial arrhythmia (N/Y)** | **3.68 (0.03, 7.33)** | **0.06** | **0.048** | N/A | N/A | N/A |
| CAD (N/Y) | -1.91 (-5.25, 1.42) | 0.02 | 0.26 | -2.25 (-5.73, 1.23) | 0.03 | 0.20 |
| HF (N/Y) | -0.85 (-4.60, 2.90) | 0.003 | 0.65 | -3.04 (-7.21, 1.12) | 0.04 | 0.15 |
| COPD (N/Y) | -2.37 (-6.01, 1.27) | 0.03 | 0.20 | -2.26 (-6.46, 1.94) | 0.02 | 0.29 |
| OSA (N/Y) | -1.04 (-2.79, 4.86) | 0.004 | 0.59 | -0.81 (-5.40, 3.77) | 0.003 | 0.72 |
| Anxiety/Depression/PTSD | 1.08 (-2.42, 4.59) | 0.006 | 0.54 | 1.26 (-2.33, 4.85) | 0.01 | 0.48 |
| Other cancer (N/Y) | 0.73 (-2.55, 4.00) | 0.003 | 0.66 | -0.17 (-3.71, 3.37) | 0.00 | 0.92 |
| Beta-blocker (N/Y) | 0.65 (-2.60, 3.90) | 0.002 | 0.69 | -1.66 (-5.20, 1.88) | 0.02 | 0.35 |
| ACE-I/ARB (N/Y) | -0.71 (-4.02, 2.60) | 0.003 | 0.67 | -3.38 (-6.86, 0.09) | 0.07 | 0.06 |
| FEV_1_, % predicted | -0.007 (-0.07, 0.06) | 0.001 | 0.83 | 0.01 (-0.06, 0.08) | 0.001 | 0.83 |
| TLC, % predicted | -0.001 (-0.08, 0.08) | 0.00 | 0.98 | -0.04 (-0.12, 0.04) | 0.02 | 0.34 |
| DL_CO_, % predicted | -0.04 (-0.11, 0.02) | 0.03 | 0.16 | -0.02 (-0.09, 0.05) | 0.01 | 0.53 |
| Tumor size, cm | 0.07 (-1.08, 1.22) | 0.00 | 0.90 | -0.28 (-1.42, 0.85) | 0.01 | 0.62 |
| Stage IA (N/Y) | 1.26 (-2.16, 4.68) | 0.008 | 0.46 | -0.45 (-4.02, 3.12) | 0.001 | 0.80 |
| Adenocarcinoma (N/Y) | 1.15 (-2.07, 4.38) | 0.008 | 0.48 | 0.94 (-2.53, 4.40) | 0.01 | 0.59 |
| Primary treatment | F-statistics | 0.04 | 0.84 | F-statistics | 0.01 | 0.78 |
| **Pre-6MWT HR, beat/min** | **0.16 (0.05, 0.28)** | **0.11** | **0.01** | 0.09 (-0.03, 0.22) | 0.05 | 0.12 |
| Pre-6MWT SBP, mmHg | 0.04 (-0.06, 0.12) | 0.009 | 0.44 | -0.01 (-0.12, 0.09) | 0.002 | 0.78 |
| Pre-6MWT DBP, mmHg | -0.05 (-0.18, 0.09) | 0.006 | 0.52 | -0.11 (-0.24, 0.03) | 0.05 | 0.13 |
| Pre-6MWT O_2_ sat, % | -0.34 (-1.15, 0.47) | 0.01 | 0.40 | -0.50 (-1.33, 0.33) | 0.03 | 0.23 |
| **HR change, beats** | **-0.40 (-0.50, -0.29)** | **0.45** | **<0.001** | **-0.42 (-0.55, -0.28)** | **0.44** | **<0.001** |
| SBP change, mmHg | -0.08 (-0.18, 0.01) | 0.04 | 0.09 | -0.09 (-0.18, 0.004) | 0.07 | 0.06 |
| DBP change, mmHg | -0.14 (-0.38, 0.10) | 0.02 | 0.26 | 0.01 (-0.25, 0.27) | 0.00 | 0.95 |
| O_2_ sat change, % | 0.12 (-0.24, 0.48) | 0.007 | 0.50 | 0.20 (-0.17, 0.57) | 0.02 | 0.29 |
| **6MWD, m** | **-0.02 (-0.03, -0.004)** | **0.10** | **0.01** | **-0.02 (-0.04, -0.01)** | **0.18** | **<0.01** |

***Bolded*** *variables indicate statistically significant associations at p <0.05.*

6MWD = six-minute walk distance; 6MWT = six-minute walk test; ACE-I = angiotensin converting enzyme inhibitor; ARB = angiotensin II receptor blocker; BMI = body-mass index; CAD = coronary artery disease; CI = confidence interval; COPD = chronic obstructive pulmonary disease; DBP = diastolic blood pressure; DL_CO_ = diffusion capacity of the lung for carbon monoxide; DM = diabetes mellitus; FEV_1_ = forced expiratory volume in 1 second; FVC = forced vital capacity; HF = heart failure; HR = heart rate; HRR = heart rate recovery; O_2_ sat = oxygen saturation; OSA = obstructive sleep apnea; PTSD = posttraumatic stress disorder; SBP = systolic blood pressure; SD = standard deviation; TLC = total lung capacity; UVA = univariable linear regression analysis

E-Table 2: Comparison of Clinical Characteristics in Participants With (N=41) and Without (N=22) ECG

| Characteristic | Delta (SE) | P-value*^*^* |
| --- | --- | --- |
| Age | -3.2 (2.0) | 0.12 |
| White race | N/A | 0.07 |
| Sex | N/A | 0.64 |
| BMI | -0.03 (1.2) | 0.98 |
| Pack years | 0.87 (8.0) | 0.91 |
| Hypertension | N/A | 0.10 |
| Hyperlipidemia | N/A | 0.75 |
| DM | N/A | 1.00 |
| CAD | N/A | 0.13 |
| HF | N/A | 0.58 |
| COPD | N/A | 1.00 |
| OSA | N/A | 0.25 |
| Psychiatric illness | N/A | 0.44 |
| Other cancer | N/A | 0.62 |
| Beta-blocker | N/A | 0.81 |
| ACE-I/ARB | N/A | 1.00 |
| FEV_1_/FVC, % | 4.7 (3.6) | 0.20 |
| FEV_1_, % predicted | 1.2 (6.2) | 0.85 |
| TLC, % predicted | -6.8 (5.9) | 0.26 |
| DL_CO_, % predicted | 2.9 (6.3) | 0.65 |
| Surgical resection | N/A | 0.47 |
| Pre-6MWT HR | 3.8 (3.3) | 0.26 |
| Pre-6MWT SBP | -7.1 (4.5) | 0.12 |
| Pre-6MWT DBP | -2.6 (3.0) | 0.38 |
| Pre-6MWT O_2_ sat | -0.23 (0.5) | 0.65 |
| HR change | -0.3 (2.8) | 0.93 |
| SBP change | 0.08 (4.3) | 0.99 |
| **DBP change** | **4.4 (1.6)** | **0.01** |
| O_2_ sat change | -0.4 (1.1) | 0.75 |
| 6MWD | 38.8 (30.1) | 0.20 |
| HRR | -2.6 (1.6) | 0.11 |

*^*^Paired-sample t-tests for continuous and Fisher’s exact test for categorical variables.*

***Bolded*** *variables indicate statistically significant difference at p <0.05.*

6MWD = six-minute walk distance; 6MWT = six-minute walk test; ACE-I = angiotensin converting enzyme inhibitor; ARB = angiotensin II receptor blocker; BMI = body-mass index; CAD = coronary artery disease; COPD = chronic obstructive pulmonary disease; DBP = diastolic blood pressure; DL_CO_ = diffusion capacity of the lung for carbon monoxide; DM = diabetes mellitus; ECG = electrocardiogram; FEV_1_ = forced expiratory volume in 1 second; FVC = forced vital capacity; HF = heart failure; HR= heart rate; HRR = heart rate recovery; O_2_ sat = oxygen saturation; OSA = obstructive sleep apnea; SBP = systolic blood pressure; SE = standard error; TLC = total lung capacity

E-Table 3: UVA – Determinants of HRV

|  | Ln-SDNN (N = 41) | | | Ln-rMSSD (N = 41) | | |
| --- | --- | --- | --- | --- | --- | --- |
| Variable | β | R^2^ | P-value | β | R^2^ | P-value |
| **Age, per year** | **0.03** | **0.11** | **0.04** | 0.02 | 0.09 | 0.052 |
| White race (N/Y) | 0.05 | 0.001 | 0.87 | 0.03 | 0.00 | 0.90 |
| Sex (F/M) | 0.23 | 0.01 | 0.63 | 0.21 | 0.01 | 0.64 |
| BMI, per kg/m^2^ | 0.01 | 0.01 | 0.48 | 0.02 | 0.02 | 0.42 |
| Active smoker (N/Y) | 0.09 | 0.01 | 0.67 | 0.06 | 0.002 | 0.76 |
| Hypertension (N/Y) | -0.22 | 0.02 | 0.36 | -0.21 | 0.02 | 0.36 |
| Hyperlipidemia (N/Y) | 0.20 | 0.02 | 0.43 | 0.20 | 0.02 | 0.43 |
| DM (N/Y) | 0.28 | 0.04 | 0.22 | 0.25 | 0.03 | 0.26 |
| CAD (N/Y) | -0.05 | 0.001 | 0.83 | -0.03 | 0.001 | 0.88 |
| HF (N/Y) | -0.14 | 0.01 | 0.58 | -0.15 | 0.01 | 0.53 |
| COPD (N/Y) | 0.29 | 0.04 | 0.21 | -0.26 | 0.04 | 0.24 |
| OSA (N/Y) | 0.26 | 0.03 | 0.25 | 0.24 | 0.03 | 0.27 |
| Psychiatric illness (N/Y) | -0.15 | 0.01 | 0.53 | -0.14 | 0.01 | 0.54 |
| Other cancer (N/Y) | -0.23 | 0.03 | 0.26 | -0.22 | 0.03 | 0.28 |
| Beta-blockers (N/Y) | 0.17 | 0.02 | 0.39 | 0.17 | 0.02 | 0.40 |
| ACE-I/ARB (N/Y) | -0.14 | 0.01 | 0.49 | -0.14 | 0.01 | 0.49 |
| FEV_1_, % predicted | -0.004 | 0.02 | 0.37 | -0.004 | 0.02 | 0.35 |
| TLC, % predicted | 0.01 | 0.11 | 0.053 | 0.01 | 0.11 | 0.06 |
| DL_CO_, % predicted | -0.002 | 0.01 | 0.57 | -0.002 | 0.01 | 0.57 |
| Pre-6MWT HR, beats/min | -0.01 | 0.08 | 0.08 | -0.01 | 0.08 | 0.08 |
| Pre-6MWT SBP, mmHg | 0.003 | 0.01 | 0.67 | 0.003 | 0.01 | 0.60 |
| Pre-6MWT DBP, mmHg | -0.001 | 0.00 | 0.91 | -0.001 | 0.00 | 0.95 |
| Pre-6MWT O_2_ sat, % | 0.04 | 0.02 | 0.41 | 0.04 | 0.02 | 0.45 |
| **HR change, beats/min** | **0.02** | **0.11** | **0.04** | **0.02** | **0.12** | **0.03** |
| SBP change, mmHg | 0.01 | 0.02 | 0.38 | 0.01 | 0.02 | 0.36 |
| DBP change, mmHg | -0.02 | 0.04 | 0.21 | -0.02 | 0.04 | 0.22 |
| O_2_ sat change, % | -0.03 | 0.05 | 0.16 | -0.03 | 0.06 | 0.13 |
| **HRR, beats** | **-0.04** | **0.15** | **0.01** | **-0.04** | **0.16** | **0.01** |
| 6MWD, m | 0.0001 | 0.00 | 0.93 | 0.00 | 0.001 | 0.84 |

***Bolded*** *variables indicate statistically significant associations at p < 0.05.*

6MWD = six-minute walk distance; 6MWT = six-minute walk test; ACE-I = angiotensin converting enzyme inhibitor; ARB = angiotensin II receptor blocker; BMI = body-mass index; CAD = coronary artery disease; COPD = chronic obstructive pulmonary disease; DBP = diastolic blood pressure; DL_CO_ = diffusion capacity of the lung for carbon monoxide; DM = diabetes mellitus; FEV_1_ = forced expiratory volume in 1 second; FVC = forced vital capacity; HF = heart failure; HR = heart rate; HRR = heart rate recovery; HRV = heart rate variability; Ln = natural logarithm; O_2_ sat = oxygen saturation; OSA = obstructive sleep apnea; PTSD = posttraumatic stress disorder; SBP = systolic blood pressure; SD = standard deviation; SDNN = standard deviation of normal-to-normal R-R intervals; rMSSD = root mean square of successive differences; TLC = total lung capacity; UVA = univariable linear regression analysis
